# Moderate episodic prenatal alcohol does not impact female offspring fertility in rats

**DOI:** 10.1101/2020.01.21.914648

**Authors:** Elizabeth K McReight, Seng H Liew, Sarah E Steane, Karla J Hutt, Karen M Moritz, Lisa K Akison

## Abstract

Prenatal alcohol exposure (PAE) has been associated with reproductive dysfunction in offspring. However, studies in females, particularly examining long-term infertility or impacts on ovarian reserve, are lacking. The current study utilised a moderate, episodic exposure model in rats to mimic ‘special occasion’ drinking, which is reported to be common during pregnancy. Our objective was to examine the consequences of this prenatal alcohol exposure on reproductive parameters in female offspring. Pregnant Sprague Dawley rats were treated with either an ethanol gavage (1g EtOH/kg body weight), or an equivalent volume of saline, on embryonic days 13.5 and 14.5 of pregnancy, resulting in a peak blood alcohol concentration of ∼0.04%. Neonatal female offspring were examined for molecular markers regulating early follicle numbers in the ovary and unbiased stereology used to quantify primordial and early growing follicle numbers. Puberty onset (age at vaginal opening and first estrous) was measured post-weaning and estrous cycles, reproductive hormones (progesterone and estradiol) and pregnancy success measured in adults (5-6 months of age). We found no evidence that any of these reproductive parameters were significantly altered by PAE in this model. This animal study provides some reassurance for women who may have consumed a small amount of alcohol during their pregnancy. However, previously published effects on offspring metabolism using this model reinforce avoidance of alcohol during pregnancy.

## Introduction

Women are increasingly deferring childbirth until later in life (Alviggi, et al. 2009), but despite recent advances in assisted reproductive technologies, there are no strategies to address declining fertility with age. Unlike men, women are born with their lifetime supply of gametes that are established before birth (Zuckerman 1951). These are contained within non-growing primordial follicles, defined as the ovarian reserve (Findlay, et al. 2015b), and declines exponentially with age as the pool of oocytes are depleted by ovulation or, more commonly, by atresia (Finch 2014). However, the number initially established varies widely between individuals, with a 70-fold range predicted at birth corresponding with a predicted age range at menopause of ∼38-60 years (Wallace and Kelsey 2010). The putative molecular factors involved in the establishment and maintenance of the ovarian reserve are just beginning to be elucidated (Findlay, et al. 2015a, Kelsey, et al. 2012, Pangas 2012), with a balance between factors involved in follicle quiescence, activation or apoptosis integral to ensuring the longevity of the reproductive lifespan (Reddy, et al. 2010).

Importantly, environmental and nutritional exposures during the prenatal establishment of ovarian reserve have been identified as critical determinants for the length of the female reproductive lifespan and subsequent fertility, especially later in life (Richardson, et al. 2014). For example, animal models show that maternal exposure to endocrine disrupting chemicals such as the plasticiser bisphenol-A (BPA) (Hunt, et al. 2012), cigarette smoking (Camlin, et al. 2016) and malnutrition (Bernal, et al. 2010, Mossa, et al. 2013, Winship, et al. 2018) can all impact on the endocrine and nutritional milieu during pregnancy, resulting in reduced follicle numbers. However, a common maternal insult that has not received as much attention is prenatal alcohol exposure (PAE). Most health authorities advise abstinence from alcohol consumption while pregnant or planning a pregnancy (National Health and Medical Research Council 2009, World Health Organisation 2004). Despite this, recent reports suggest that women are consuming alcohol both prior to pregnancy recognition (Ishitsuka, et al. 2019, McCormack, et al. 2017, Muggli, et al. 2016) and late in pregnancy (Muggli, et al. 2016, Umer, et al. 2020), with rates as high as 50-60% in these studies and ∼10% globally (Popova, et al. 2017).

Although the neurological and behavioural deficits in offspring arising from PAE are well known, emerging evidence suggests that a much broader range of body systems can be affected. This includes the reproductive system, with a recent systematic review identifying deficits in males and females from both clinical and preclinical studies (Akison, et al. 2019). However, this review identified a general paucity of studies examining the female reproductive system (3 clinical and 12 preclinical), with almost all studies focussing on age at first menarche/puberty onset, which was significantly delayed in PAE offspring compared to controls in most studies (Boggan, et al. 1979, Esquifino, et al. 1986, McGivern, et al. 1992, McGivern and Yellon 1992, Robe, et al. 1979). Importantly, no studies to-date have examined impacts of PAE on ovarian reserve or long-term fertility. There are a few studies reporting direct effects of alcohol consumption on ovarian reserve, with one study reporting heavy binge drinking impacted on plasma AMH levels, a proxy measure for ovarian reserve, in a group of African American women (Hawkins Bressler, et al. 2016), and evidence of earlier age at menopause in women with alcohol use disorder (Choi, et al. 2017, Gavaler 1985).

Other direct effects of alcohol on the female endocrine system have been reported (Rachdaoui and Sarkar 2017), resulting in irregular menstrual/estrous cycles, anovulation, early menopause and elevated estradiol (E2) levels in both women and rodents. High alcohol consumption has also been suggested to negatively impact on assisted reproductive outcomes, with reduced oocytes retrieved, lower rates of pregnancy and increased risk for miscarriage (see Van Heertum and Rossi (2017) for review).

Previous studies of PAE on female reproductive outcomes in offspring have typically been conducted in women with very high levels of alcohol use (clinical) or using chronic, high doses (preclinical) (see Akison, et al. (2019) for review). However, this is not representative of typical reported drinking behaviours in pregnant women, which are often episodic and at low-moderate levels of alcohol, referred to as ‘special occasion’ drinking (McCormack, et al. 2017, Muggli, et al. 2016). This study, using a rat model of acute, low-level exposure that has been previously reported to result in a blood alcohol concentration (BAC) of ∼0.05%, did not result in fetal growth restriction but still produced sex-specific adverse metabolic outcomes in adult offspring (Nguyen, et al. 2019). The timing of exposure corresponded with germ cell cyst or ‘nest’ formation in the ovaries of female offspring, with breakdown of these nests and encapsulation of individual oocytes by somatic cells to form primordial follicles occurring in late gestation and peaking in early postnatal life in rodents (Sarraj and Drummond 2012).

Our objective was to examine the impact of this PAE model on the establishment of offspring ovarian reserve and subsequent impacts on puberty onset in adolescence and fertility parameters in adulthood. We hypothesised that PAE would reduce the ovarian reserve in neonates, delay puberty onset, disrupt estrous cycling in adults and reduce fertility in adult female offspring.

## Materials and methods

### Animal model

Ethics approval for all animal experimentation was obtained from the University of Queensland Anatomical Biosciences Animal Ethics Committee (SBMS/AIBN/521/15/NHMRC) prior to commencement of the study and was conducted in accordance with the Australian Code for the Care and Use of Animals for Scientific Purposes (2013, 8th Edition). Reporting of animal experiments conforms to the ARRIVE guidelines (Kilkenny and Altman 2010, Kilkenny, et al. 2010).

A detailed description of the source of animals, housing conditions and treatments has been previously reported (Nguyen, et al. 2019). Briefly, outbred, nulliparous Sprague-Dawley rats were housed in a temperature- and humidity-controlled environment with an artificial 12 h reversed light-dark cycle and provided with standard laboratory rat chow (Rat & Mouse Meat-Free Diet, Specialty Feeds, Glen Forrest WA, Australia) and water ad libitum. After an initial acclimation period, dams were mated with a proven stud male (1200-1700) and pregnancy confirmed via presence of a seminal plug. The following morning was designated as embryonic day (E) 0.5. Once pregnant, dams were randomly allocated to either receive ethanol (EtOH) or saline (Control) via oral gavage at E13.5 and E14.5. EtOH treated females (*n* = 16) received 18% v/v EtOH in saline solution (0.9% NaCl) at a dose of 1 g/kg body weight, while Control females (*n* = 14) received an equivalent volume of saline. Water and chow consumption were measured daily from E12.5 (one day prior to first gavage) and weight gain monitored throughout pregnancy. Day of birth was designated postnatal day (PN) 0. Dams and offspring were allocated either to neonatal or adolescent/adult studies of reproductive parameters. Figure 1 provides a flow chart showing the dams allocated to each arm of the study and the number of offspring that contribute to each analysis at each age. Only 1-2 females were used from each litter for each experiment to remove potential litter effects, as recommended for DOHaD studies by Dickinson, et al. (2016). The exception was monitoring puberty onset, where litter averages were obtained across 5-7 offspring per litter.

**Figure 1.**
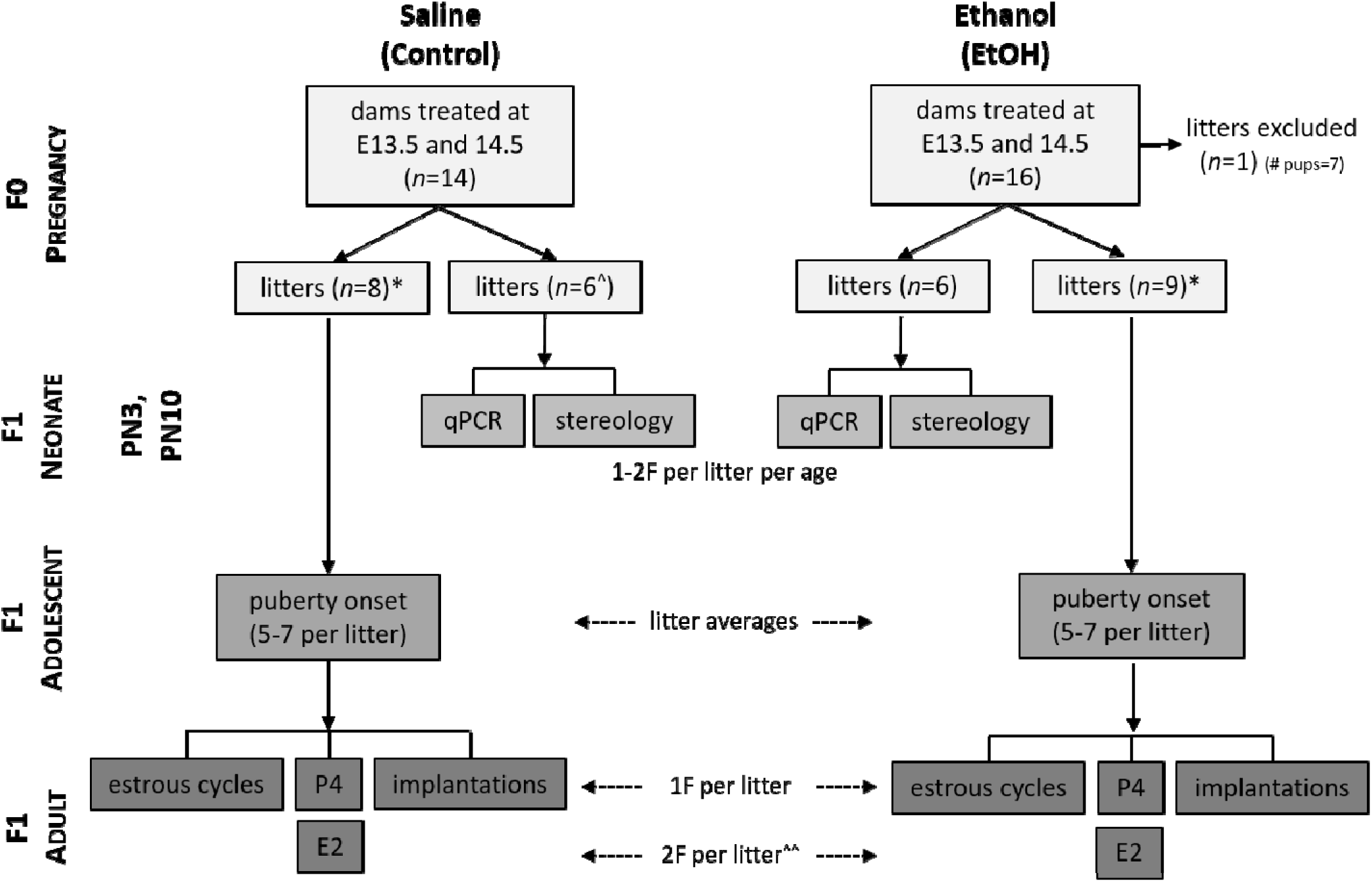
Flow chart of animals treated and offspring used in each experiment. Pregnant dams were treated with either saline or 1 g/kg body weight ethanol (EtOH) via gavage. Note that one EtOH litter was excluded from further experiments given the litter size was 7 (mean, range for other litters: 15, 11-19). * A separate cohort of offspring from these litters were also examined for metabolic outcomes (reported in Nguyen et al. 2019). ^ *n*=5 at PN3 due to aging error for 1 litter. ^^ Estradiol measured in a separate group of females to those used for estrous cycles etc. E, embryonic day; E2, estradiol; F, female; P4, progesterone; PN, postnatal day; qPCR, quantitative real-time polymerase chain reaction.

### Blood and tissue collection

Approximately 150 μl whole blood was collected from pregnant dams via a tail tip bleed at 1 h and 5 h post gavage on E13.5 and 14.5 to measure blood alcohol concentration (BAC) (Nguyen, et al. 2019). An additional tail tip bleed at E19.5 was used to measure plasma progesterone levels. Female offspring from neonatal cohort litters were collected at PN3 and PN10 (1 – 2 per litter per time point), weighed, culled via decapitation and ovaries collected via microdissection. Ovaries were either snap frozen using liquid nitrogen and stored at - 80°C for molecular analysis or fixed in 4% paraformaldehyde (PFA) for histological analysis. At 6 months of age, two female offspring from adult cohort litters were culled via carbon dioxide asphyxiation at proestrous. Whole blood was collected into EDTA tubes via cardiac puncture for hormone analysis (estradiol). An additional female from each litter was used for mating experiments as described below and whole blood collected via cardiac puncture at day 9.5-11.5 of pregnancy for hormone analysis (progesterone). All blood samples were immediately centrifuged at 4000 rpm for 10 min at 4°C and plasma separated, aliquoted and stored at -20°C until subsequent analysis.

### Measurement of blood alcohol concentration (BAC) and steroid hormones

Plasma BAC was determined by Pathology Queensland (Queensland Health) using an alcohol dehydrogenase enzymatic assay (Beckman Coulter, Lane Cove, NSW, Australia; reference #474947) as previously reported (Nguyen, et al. 2019). Steroid hormones were analysed by radioimmunoassay (RIA) as previously described (Kalisch-Smith, et al. 2019). Briefly, plasma estradiol was analysed by ultra-sensitive RIA (Beckman Coulter; reference # DSL-4800), with a limit of detection of 22.4 pmol/L and intra-assay coefficient of variation (CV) of 1.6%. Plasma progesterone was analysed using an RIA developed in-house and progesterone antiserum C-9817 (Bioquest, North Ryde, NSW, Australia), with a limit of detection of 0.32 nmol/L and an intra-assay CV of 2.7%.

### Assessment of puberty onset and first estrous in adolescent offspring

From PN30, female offspring were monitored for puberty onset, characterised by the complete loss of the vaginal membrane, as previously described (McGivern and Yellon 1992). Following puberty onset, each animal was monitored daily using a mouse-sized vaginal impedance probe and the EC40 estrous cycle monitor (Fine Science Tools) to detect electrical impedance >4.0 k Ω, indicating the first pro-estrous.

### Estrous cycle tracking and fertility assessments in adult offspring

At six months of age, one female from each litter had their estrous cycles monitored daily using the EC40 estrous cycle monitor (Fine Science Tools) as described above for 21 days. Following this monitoring period, the same animals were mated at pro-estrous with a proven stud male (1200-1700) and pregnancy confirmed via presence of a seminal plug. The following morning was designated as embryonic day (E) 0.5. Once pregnant, dams were housed singly. At E9.5-11.5, pregnant offspring were culled by CO_2_ asphyxiation, cardiac blood was collected into EDTA tubes for future hormone analysis and the uterus was removed. Implantation and reabsorption sites were counted as a measure of fertility.

Implantations rather than live birth rate was used to assess fertility to minimise animal wastage, although we acknowledge that potential losses in late pregnancy may have been missed using this method.

### RNA extraction and quantitative PCR (qPCR)

RNA was extracted from frozen neonatal ovarian tissue (both ovaries from 1-2 animals per litter, *n* = 10-12 per time-point per group; see Supplemental Table 1 for more details) using either: PN3 ovaries - QIAzol Lysis Reagent (Qiagen, Chadstone, VIC, Australia) and Glycoblue coprecipitant (Thermo Fischer Scientific, Richlands, QLD, Australia) to assist with pellet visualisation as per the manufacturer’s instructions, with an additional overnight precipitation at -20°C; or PN10 ovaries - QIAGEN RNeasy Minikit (Qiagen) according to the manufacturer’s instructions. All ovaries were initially homogenised using a 20 gauge needle and 1 mL syringe to assist with lysis. RNA concentration was quantified using a Nanodrop 2000 spectrophotometer (Thermo Fischer Scientific) and yields were typically ≥25 ng/µL and ≥ 130 ng/µL of total RNA for PN3 and PN10 ovaries respectively. The 260/280 ratio for all ≥ samples was ≅ 2.0. cDNA synthesis was performed using the iScript Reverse Transcription (RT) Supermix (Bio-Rad Laboratories, Gladesville, NSW, Australia), according to the manufacturer’s instructions. Each reaction contained 200 ng of RNA, reverse transcribed to produce 20 ng/µL cDNA. RT reactions were performed on a PCR Express Thermal Cycler (Thermo Fisher Scientific). Samples were then diluted 1:10 for a working concentration of 2 ng/ µL and stored at -20°C.

Expression of genes involved in regulation of primordial follicle numbers were analysed using real-time quantitative polymerase chain reaction (qPCR) (see Table 1 for details of specific genes). qPCR reactions were performed in duplicate on an Applied Biosystems Quantstudio 6 Flex Real-Time PCR System (ThermoFisher Scientific) using 4 ng cDNA, Taq PCR Master Mix (Qiagen; catalogue #201443) and Taqman Assay-on-Demand primer/probe sets (Thermo Fischer Scientific; see Table 1 for details) per 10 µL reaction. Relative gene expression was determined using the comparative threshold method (ΔΔCT) and normalised to the mean of *Hprt* run in duplicate as the endogenous control. Only one endogenous control gene was found to be stably expressed across all samples, regardless of experimental group, from a previous pilot study of 10 commonly used endogenous control genes in similar neonatal ovary samples exposed to prenatal alcohol (data not shown; see footnote to Table 1 for other genes tested). Fold-change was expressed relative to the average of the saline control group at each age.

**Table 1.** Primers used for real-time quantitative PCR analysis of genes regulating primordial follicle numbers. All primers were Assay-on-Demand primer/probe sets from Thermo Fisher Scientific (Richlands, QLD, Australia).

| Gene name | Gene symbol | Assay ID | Amplicon size (bp) | Accession number(s) |
| --- | --- | --- | --- | --- |
| BCL-2 associated X protein | *Bax* | Rn01480161_g1 | 63 | NM_017059.2 |
| BCL-2 antagonist/killer 1 | *Bak1* | Rn00587491_m1 | 64 | NM_053812.1<br>XM_006256101.2 |
| B cell leukemia/lymphoma 2 | *Bcl2* | Rn99999125_m1 | 104 | NM_016993.1 |
| BCL2-like 1 | *Bcl2l1* | Rn00437783_m1 | 65 | NM_001033670.1<br>XM_006235265.2 |
| BCL2-like 11 (apoptosis facilitator) | *Bcl2l11* | Rn00674175_m1 | 64 | NM_022612.1<br>NM_171988.2<br>NM_171989.1<br>XM_006234983.2<br>XM_006234985.2<br>XM_006234987.2 |
| BCL2 binding component 3 (p53-upregulated modulator of apoptosis) | *Bbc3 (Puma)* | Rn00597992_m1 | 62 | NM_173837.2 |
| Inhibin alpha | *Inha* | Rn00561423_m1 | 58 | NM_012590.2 |
| Anti-mullerian hormone | *Amh* | Rn00563731_g1 | 92 | NM_037034.1 |
| Serine/threonine kinase 11 (Liver kinase B1) | *Stk11 (Lkb1)* | Rn01535544_m1 | 59 | NM_001108069.1<br>XM_006240910.1<br>XM_008765096.1 |
| Chemokine (C-X-C motif) ligand 12 | *Cxcl12* | Rn00573260_m1 | 60 | NM_001033882.1<br>NM_001033883.1<br>NM_022177.3 |
| Chemokine (C-X-C motif) receptor 4 | *Cxcr4* | Rn00573522_s1 | 58 | NM_022205.3 |
| Phosphatase and tensin homolog | *Pten* | Rn00477208_m1 | 73 | NM_031606.1 |
| Hypoxanthine phosphoribosyltransferase * | *Hprt* | Rn01527840_m1 | 64 | NM_012583.2 |

\* Endogenous control. Note that a previous (unpublished) pilot study, using neonatal ovaries collected at similar ages from female offspring exposed to prenatal alcohol, found that *Hprt* was the only stably expressed gene across EtOH-treated and control ovaries from 10 tested (*Actb*, *Rpl19*, *Hprt*, *Mrpl1*, *Pgk1*, *Sdha*, *Rpl0*, *Rpl13*, *Ppia*, *18S*).

### Ovarian histology and stereology

Ovarian follicle counts were performed on ovaries collected from neonatal offspring at PN3 and PN10 using unbiased stereology, considered the ‘gold standard’ for quantification of cells in tissue sections (Geuna and Herrera-Rincon 2015). One ovary from each neonate (1-2 animals per litter, *n* = 8-10 per time-point per group; see Supplemental Table 1 for more details) was fixed in 4% PFA, processed manually via increasing concentrations of EtOH and xylene, and embedded in paraffin wax. Ovaries were serially sectioned at 5µm and every 9^th^ section stained with Period Acid Schiff (PAS) and counterstained with haematoxylin.

Primordial, transitional and primary follicles were classified as previously described (Myers, et al. 2004). Briefly, single oocytes surrounded by a complete or incomplete layer of squamous granulosa cells were considered non-growing, primordial follicles (i.e. the ovarian reserve); single oocytes surrounded by a single layer of cuboidal granulosa cells were considered primary follicles; and a mixture of squamous and cuboidal granulosa cells were considered transitional follicles. Where oocytes were still clustered together in an ovarian cyst or ‘nest’ prior to follicle formation (at PN3 only), these were counted separately.

The optical disector/fractionator method was used to quantify primordial, transitional and primary follicles as previously described (Myers, et al. 2004, Stringer, et al. 2019). All counts were performed blinded to treatment/control. Briefly, every ninth section (i.e. f1 = sampling fraction = 1/9) was examined on a Nikon Upright Stereology and Slide Scanning microscope (SciTech Pty Ltd, Preston, VIC, Australia) with motorised stage at 40X magnification using the Stereo Investigator stereology system (Version 2018, MBF Bioscience, Williston, VT, USA). Using the Stereo Investigator software, a sampling grid (grid size 200 µm x 200 µm = 40, 000 µm^2^) was overlaid over the section, with a 95 µm x 95 µm (9025 µm^2^) counting frame overlaid in each sampling grid square (f2 = 9025/40, 000). Each counting frame consisted of inclusion and exclusion boundaries, with oocytes exhibiting clearly defined nuclei, within the counting frame or on the inclusion boundary, counted to provide raw counts of each follicle type (Q-). As thin sections were used, the entire 5 µm was counted (f3 = 5/5). To estimate the total number of each follicle type per ovary (N), the following equation was used: N = Q-_(follicle)_ x (1/f1) x (1/f2) x (1/f3).

Total secondary and antral follicle numbers were estimated per ovary by counting follicles with a clearly defined nuclei within the oocyte in every 36^th^ section at 10X magnification. Growing follicles were classified as previously described (Myers, et al. 2004). Briefly, secondary follicles were characterised by more than one layer of cuboidal granulosa cells surrounding the oocyte without any visible antral spaces. Antral follicles were defined by a clear antral space. Atretic follicles could not be quantified due to the difficulty of identifying apoptotic cells in paraffin sections. Raw counts were multiplied by the sampling interval (i.e. 36) to obtain total counts per ovary.

### Statistical analyses

All raw data for each analysis are provided in Supplemental Table 1. Analyses were conducted using GraphPad Prism 7.0 (GraphPad Software, San Diego, CA, USA) and data presented as mean ± SEM. Prior to hypothesis testing, data were tested for a normal distribution using the D’Agostino-Pearson test. Control and EtOH-exposed groups were compared for each parameter using a Student’s t-test (parametric data) or a Mann-Whitney U-test (non-parametric data). Where there was a significant difference in the standard deviations between groups, a Student’s t-test with Welch’s correction was conducted. Fishers exact test was used to compare the cumulative percentage of offspring in each group reaching puberty or estrous at each age. Significance level was *P* <0.05 for all statistical tests, with a Bonferroni correction for multiple testing applied to analysis of gene expression (12 genes = 0.05/12 = 0.004).

## Results

### Maternal parameters and postnatal growth

Data from dams used to produce offspring for the adolescent/adult arm of the study have been previously reported (Nguyen, et al. 2019). However, these animals were also included in the summary of maternal parameters for completeness. All dams were of similar weight at mating and at time of gavage and pregnancy weight gain was similar across both groups pre- and post-gavage (Table 2). Water and chow consumption was also not affected by treatment (Table 2). Note that Nguyen, et al. (2019) also reported that for dams specifically used to produce offspring for the adolescent/adult arm of the study, there were no significant differences in blood glucose or energy intake post-gavage. There were also no differences in pregnancy outcomes between EtOH and Control dams, including litter sex ratio, litter size and plasma progesterone levels measured in late gestation (Table 2). Nguyen, et al. (2019) also reported that for dams specifically used to produce offspring for the adolescent/adult arm of the study, there were no significant differences in the number of implantation scars, suggesting no differences in potential late pregnancy loss prior to birth. Note that one dam in the EtOH group had a litter of only 7 pups due to a suspected blockage in one uterine horn (no implantation scars were observed) and so this litter was excluded from subsequent analyses (both here and in Nguyen, et al. (2019)). BAC was measured in 6 out of 14 Control dams, with BAC below the limit of detection at each time point. In EtOH-treated dams, mean BAC was ∼42 mg/dL (∼0.04%) at 1 h following gavage at E13.5 and E14.5, but by 5 h post-gavage, was below the limit of detection on both days (Table 2).

**Table 2.** Maternal parameters for rat dams treated with saline (control) or ethanol (EtOH). All animals treated by oral gavage on embryonic day 13.5 and 14.5.

| **Parameter** | **Control**<br>(n = 14) | **EtOH**<br>(n = 15) | **P value** |
| --- | --- | --- | --- |
| Body weight at mating (g) | 252 ± 3 | 256 ± 4 | 0.41 |
| Body weight at E13.5 (g) | 309 ± 4 | 315 ± 4 | 0.30 |
| **Total weight gain (g)** |  |  |  |
| Pre-gavage (mating to E12.5) | 52 ± 3 | 56 ± 3 | 0.28 |
| Post-gavage (E15.5 to birth) | 91 ± 5 | 93 ± 3 | 0.75 |
| **Water consumption (mL/day)** |  |  |  |
| Pre-gavage (E12.5)<sup>^</sup> | 31.9 ± 1.5 | 29.3 ± 1.2 | 0.19 |
| During gavage (E13.5-E14.5) | 28.5 ± 1.6 | 26.9 ± 1.2 | 0.42 |
| Post-gavage (E15.5 to birth) | 37.9 ± 1.2 | 35.3 ± 1.2 | 0.12<sup>*</sup> |
| **Chow consumption (g/day)** |  |  |  |
| Pre-gavage (E12.5)<sup>^</sup> | 24.7 ± 0.8 | 24.9 ± 0.8 | 0.83 |
| During gavage (E13.5-E14.5) | 22.3 ± 0.9 | 21.2 ± 0.7 | 0.37 |
| Post-gavage (E15.5 to birth) | 26.2 ± 0.7 | 26.1 ± 0.6 | 0.88 |
| **Pregnancy variables** |  |  |  |
| Litter sex ratio (M:F) | 0.95 ± 0.1 | 0.99 ± 0.1 | 0.71<sup>*</sup> |
| Litter size | 14<sup>†</sup> | 15<sup>†</sup> | 0.11 |
| Plasma progesterone (nmol/L)<sup>%</sup> | 52.7 ± 5.5 | 71.8 ± 9.9 | 0.14<sup>*</sup> |
| **Blood alcohol concentration (BAC) (mg/dL)<sup>^</sup>** |  |  |  |
| 1h post E13.5 gavage | <LD<sup>‡</sup> | 41.3 ± 6.2 | - |
| 5h post E13.5 gavage | <LD<sup>‡</sup> | <LD | - |
| 1h post E14.5 gavage | <LD<sup>‡</sup> | 42.9 ± 6.3 | - |
| 5h post E14.5 gavage | <LD<sup>‡</sup> | <LD | - |

Data are presented as mean ± SEM. P values were obtained using an unpaired Student's t-test. Threshold for significance: $P < 0.05$ . <LD = below the limit of detection (5 mg/dL or 0.005%); E = embryonic day; EtOH = ethanol; M = male; F = female. <sup>\*</sup> Indicates data not normally distributed and analysed with a non-parametric Mann-Whitney U-test. <sup>^</sup> Indicates not all samples included in analysis. See Supplemental Data 1 for details. <sup>†</sup> Denotes parameters expressed as whole pups. <sup>‡</sup> Measured in 6 animals only. <sup>%</sup> Measured at E19.5; 1 sample missing from each group.

There was no difference in neonatal pup offspring weights at ovary collection on PN3 and PN10 between EtOH and Control groups (Table 3). There were also no differences in weights of adolescent offspring at PN28, one week post-weaning and just prior to monitoring for puberty onset from PN30 (Table 3). Adult offspring at 6 months of age were all of similar weight across EtOH and Control groups before measurement of reproductive parameters at this age (Table 3). Nguyen, et al. (2019) reported no difference in pup weight just after birth (PN1) for offspring used in the adolescent/adult arm of the study and no differences in weight gain up to 6 months of age.

**Table 3.**
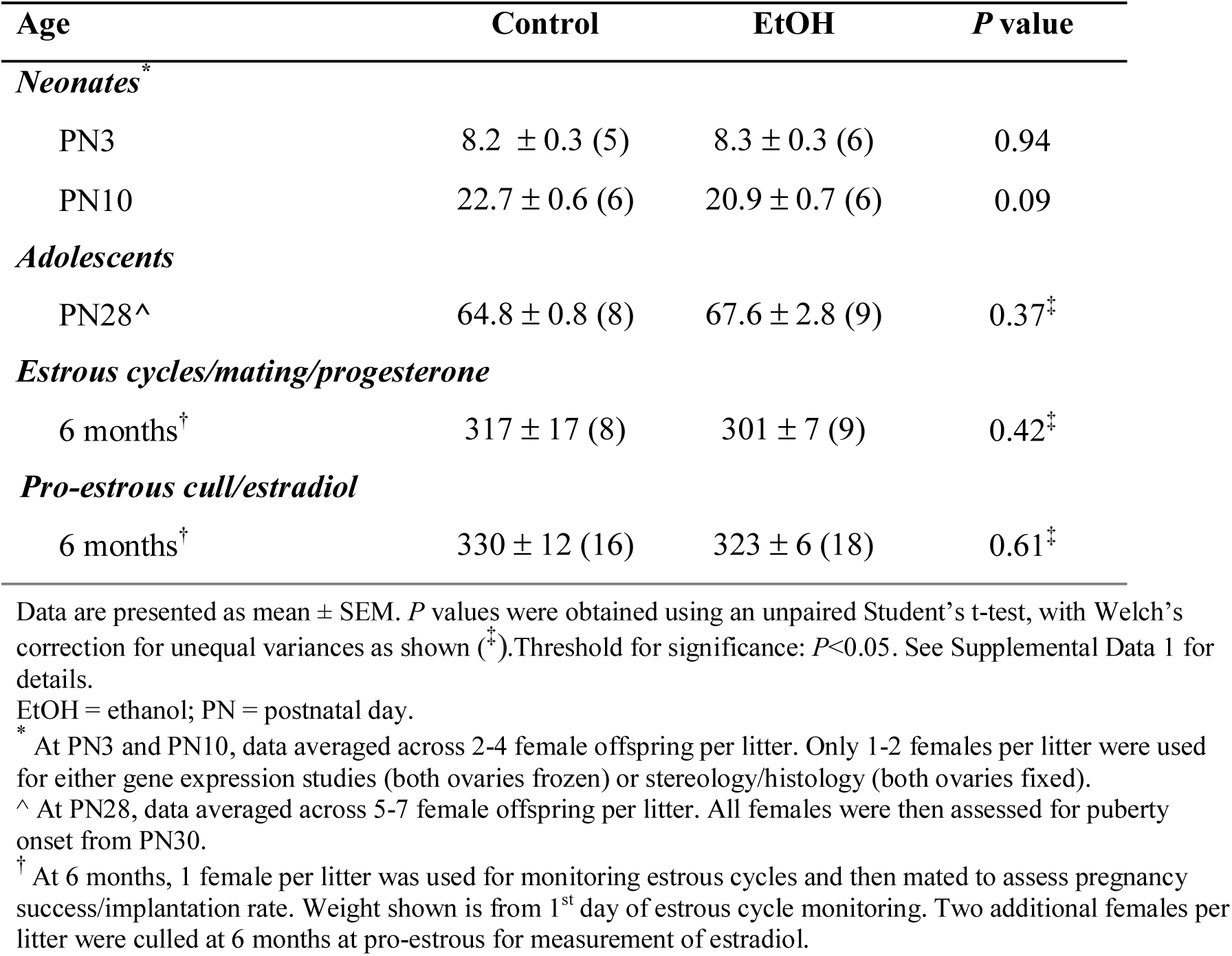
Postnatal weights (g) of female rat offspring from saline (Control) and ethanol (EtOH) treated litters used to examine reproductive parameters. All offspring from dams treated by oral gavage at embryonic day 13.5 and 14.5. Sample sizes in brackets.

| Age | Control | EtOH | P value |
| --- | --- | --- | --- |
| *Neonates*<sup>*</sup> |  |  |  |
| PN3 | 8.2 ± 0.3 (5) | 8.3 ± 0.3 (6) | 0.94 |
| PN10 | 22.7 ± 0.6 (6) | 20.9 ± 0.7 (6) | 0.09 |
| *Adolescents* |  |  |  |
| PN28<sup>^</sup> | 64.8 ± 0.8 (8) | 67.6 ± 2.8 (9) | 0.37<sup>‡</sup> |
| *Estrous cycles/mating/progesterone* |  |  |  |
| 6 months<sup>†</sup> | 317 ± 17 (8) | 301 ± 7 (9) | 0.42<sup>‡</sup> |
| *Pro-estrous cull/estradiol* |  |  |  |
| 6 months<sup>†</sup> | 330 ± 12 (16) | 323 ± 6 (18) | 0.61<sup>‡</sup> |

Data are presented as mean ± SEM. P values were obtained using an unpaired Student's t-test, with Welch's correction for unequal variances as shown (<sup>‡</sup>). Threshold for significance: P<0.05. See Supplemental Data 1 for details.
EtOH = ethanol; PN = postnatal day.
<sup>\*</sup> At PN3 and PN10, data averaged across 2-4 female offspring per litter. Only 1-2 females per litter were used for either gene expression studies (both ovaries frozen) or stereology/histology (both ovaries fixed).
<sup>^</sup> At PN28, data averaged across 5-7 female offspring per litter. All females were then assessed for puberty onset from PN30.
<sup>†</sup> At 6 months, 1 female per litter was used for monitoring estrous cycles and then mated to assess pregnancy success/implantation rate. Weight shown is from 1<sup>st</sup> day of estrous cycle monitoring. Two additional females per litter were culled at 6 months at pro-estrous for measurement of estradiol.

### Expression of factors potentially regulating follicle numbers in neonatal ovaries

PAE did not significantly impact the expression of a number of genes known to regulate primordial follicle numbers via the apoptotic pathway (*Bax, Bak1, Bcl2, Bcl2l1, Bcl2l11, Puma/Bbc3*), or positive and negative regulators of follicle growth and recruitment (*Inha, Amh, Stk11/Lkb1, Cxcl12, Pten*) at PN3 (Table 4) or PN10 (Table 5). At PN3, there was a trend for reduced expression of *Cxcr4*, involved in follicle growth, but this was at *P* = 0.04, which was not below the 0.004 threshold when adjusting for multiple testing.

**Table 4.** Expression of factors regulating primordial follicle numbers in neonatal ovaries from female rat offspring at postnatal day 3 (PN3). All offspring from dams treated by oral gavage at embryonic day 13.5 and 14.5.

| Gene of interest | Control<br>(n = 10) | EtOH<br>(n = 10) | P value |
| --- | --- | --- | --- |
| *Apoptotic pathway* |  |  |  |
| *Bax* | 1.00 ± 0.10 | 0.96 ± 0.09 | 0.79 |
| *Bak1* | 1.00 ± 0.06 | 0.91 ± 0.08 | 0.41 |
| *Bcl2* | 1.00 ± 0.11 | 0.83 ± 0.08 | 0.23 |
| *Bcl2l1* | 1.00 ± 0.07 | 0.87 ± 0.10 | 0.27 |
| *Bcl2l11* | 1.00 ± 0.11 | 0.82 ± 0.10 | 0.25 |
| *Puma/Bbc3* | 1.00 ± 0.14 | 1.26 ± 0.17<sup>^</sup> | 0.25 |
| *Follicle growth and recruitment* |  |  |  |
| *Inha* | 1.00 ± 0.07 | 0.78 ± 0.08 | 0.07 |
| *Amh* | 1.00 ± 0.20 | 0.83 ± 0.19 | 0.39<sup>*</sup> |
| *Stk11/Lkb1* | 1.00 ± 0.09 | 0.89 ± 0.11 | 0.43 |
| *Cxcl12* | 1.00 ± 0.14 | 0.90 ± 0.14 | 0.63 |
| *Cxcr4* | 1.00 ± 0.05 | 0.79 ± 0.08 | 0.04 |
| *Pten* | 1.00 ± 0.08 | 0.89 ± 0.08 | 0.34 |

Data are presented as mean ± SEM fold change, relative to *Hprt* endogenous control and normalised to the control group (i.e. set to an average fold change of 1). *P* values were obtained using an unpaired Student's t-test. Threshold for significance: *P* < 0.05 (or < 0.004, with Bonferroni correction for multiple testing). 1-2 female offspring used per litter for controls (across 5 litters) and EtOH treated (across 6 litters). See Supplemental Data 1 for details.
EtOH = ethanol.
<sup>\*</sup> Indicates data not normally distributed and analysed with a non-parametric Mann-Whitney U-test.
<sup>^</sup> 1 sample missing from this group due to qPCR error.

**Table 5.** Expression of factors regulating primordial follicle numbers in neonatal ovaries from female rat offspring at postnatal day 10 (PN10). All offspring from dams treated by oral gavage at embryonic day 13.5 and 14.5.

| Gene of interest | Control<br>(n = 10) | EtOH<br>(n = 12) | P value |
| --- | --- | --- | --- |
| *Apoptotic pathway* |  |  |  |
| *Bax* | 1.00 ± 0.08 | 0.87 ± 0.09 | 0.23* |
| *Bak1* | 1.00 ± 0.13 | 1.02 ± 0.09 | 0.28* |
| *Bcl2* | 1.00 ± 0.08 | 0.98 ± 0.09 | 0.90 |
| *Bcl2l1* | 1.00 ± 0.10 | 1.07 ± 0.10 | 0.62 |
| *Bcl2l11* | 1.00 ± 0.13 | 1.15 ± 0.12 | 0.42* |
| *Puma/Bbc3* | 1.00 ± 0.08 | 0.95 ± 0.10 | 0.69 |
| *Follicle growth and recruitment* |  |  |  |
| *Inha* | 1.00 ± 0.14 | 1.13 ± 0.13 | 0.50 |
| *Amh* | 1.00 ± 0.23 | 0.79 ± 0.09 | 0.63* |
| *Stk11/Lkb1* | 1.00 ± 0.11 | 1.04 ± 0.10 | 0.77 |
| *Cxcl12* | 1.00 ± 0.13 | 1.25 ± 0.17 | 0.27 |
| *Cxcr4* | 1.00 ± 0.11 | 1.04 ± 0.10 | 0.81 |
| *Pten* | 1.00 ± 0.10 | 1.13 ± 0.11 | 0.38* |

Data are presented as mean ± SEM fold change, relative to *Hprt* endogenous control and normalised to the control group (i.e. set to an average fold change of 1). *P* values were obtained using an unpaired Student's t-test. Threshold for significance: *P* < 0.05 (or < 0.004, with Bonferroni correction for multiple testing). 1-2 female offspring used per litter for controls (across 6 litters) and EtOH treated (across 6 litters). See Supplemental Data 1 for details.
EtOH = ethanol.
\* Indicates data not normally distributed and analysed with a non-parametric Mann-Whitney U-test.

### Ovarian reserve and follicle counts in neonatal ovaries

PAE did not impact the number of oocytes still clustered in ovarian ‘nests’ at PN3 (Figure 2A) or the number of fully-formed primordial follicles (ovarian reserve) at PN3 or PN10 (Figure 2B). At PN3, the majority (∼95%) of follicles within the ovary were primordial follicles, with ∼4% transitioning from primordial to primary and only 1% were primary follicles (Figure 2C). However, by PN10, the contribution of primordial follicles to the total follicle count within the ovary had dropped to ∼68%, with transitional (∼10%), primary (∼12%), secondary (∼9%) and even a small number of early antral (∼1%) follicles contributing to the total follicle pool (Figure 2D). There were no differences in the number of growing follicles at any stage in the ovaries from EtOH or Control neonatal offspring at PN3 or PN10 (Figure 2C-D).

**Figure 2.**
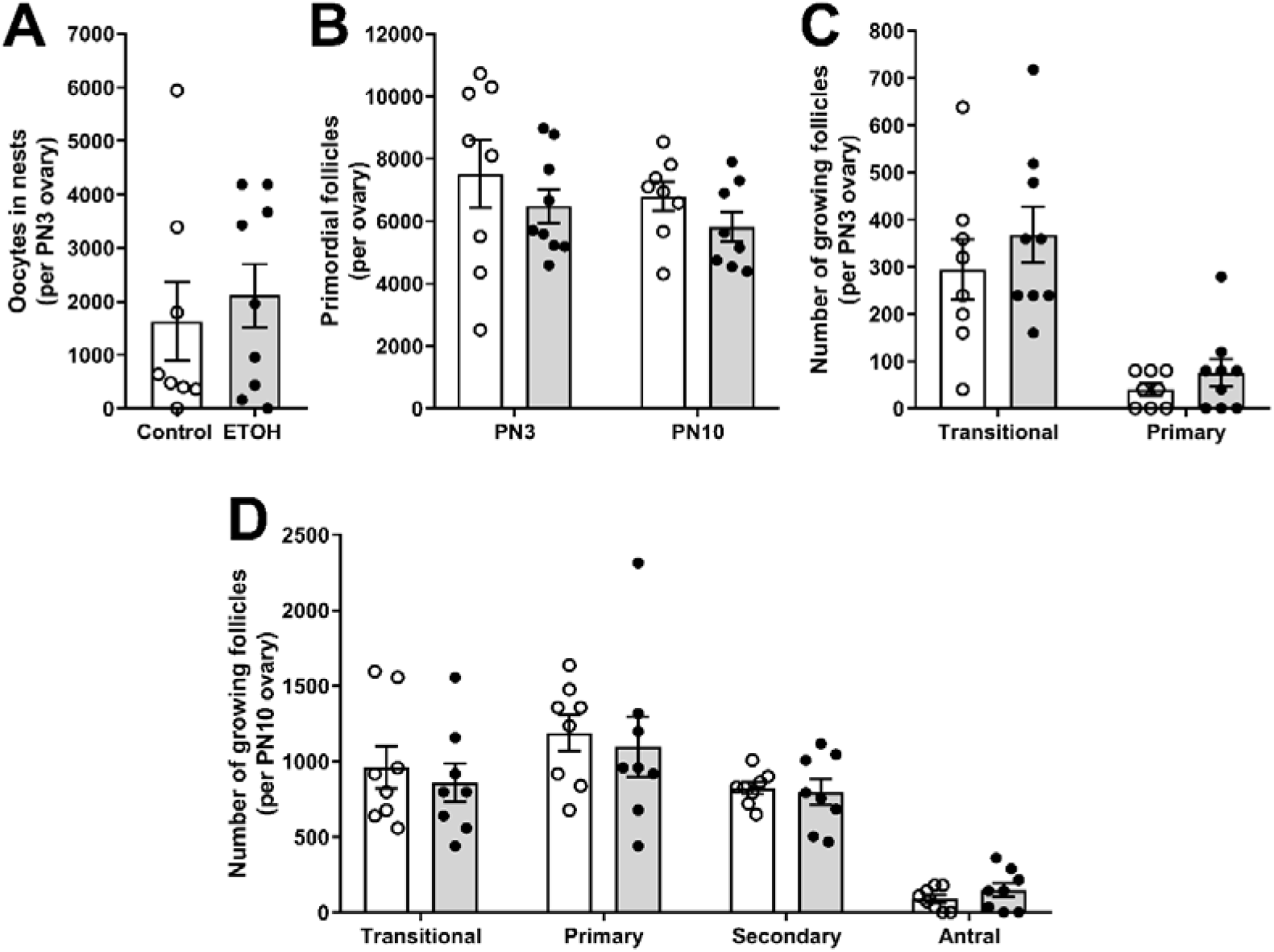
Moderate, acute prenatal alcohol exposure did not alter ovarian oocyte or follicle numbers in neonatal rat offspring. Pregnant dams were treated with ethanol (EtOH, 1 g/kg body weight) or saline (control) at embryonic day (E) 13.5 and 14.5. (A) Oocyte numbers within ovarian cysts or nests prior to follicle formation in postnatal day (PN) 3 ovaries. B) Non-growing primordial follicles (ovarian reserve) in PN3 and PN10 ovaries. (C) Early growing follicles (transitional or primary) in PN3 ovaries. D) Growing follicles (transitional, primary, secondary and antral) in PN10 ovaries. All data are presented as mean ± SEM. Open bars/circles indicate control offspring (*n* = 8 from 6 litters at both ages); grey bars/solid circles indicate EtOH offspring (*n* = 9 from 6 litters at PN3; *n* = 8 from 6 litters at PN10). Data were analysed using an unpaired t-test (parametric data) or non-parametric Mann-Whitney rank sum test to determine significant differences between groups for oocytes or each follicle type, with significance determined at *P*<0.05.

### Puberty onset and estrous cycles in adolescent offspring

PAE did not impact the timing of puberty onset in female offspring, with both the Control and EtOH groups experiencing puberty onset at an average age of PN38 (Figure 3A). The cumulative percentage of offspring achieving puberty with increasing age was similar in both groups, with all animals reaching puberty by PN45-47 (Figure 3B). Similarly, PAE did not impact the age of first estrous in female offspring, with both groups experiencing their first pro-estrous as measured by vaginal electrical impedance at an average age of PN45 (Figure 3C). The cumulative percentage of offspring achieving first estrous with increasing age was also similar in both groups, with all animals reaching this reproductive milestone by PN56-61 (Figure 3D).

**Figure 3.**
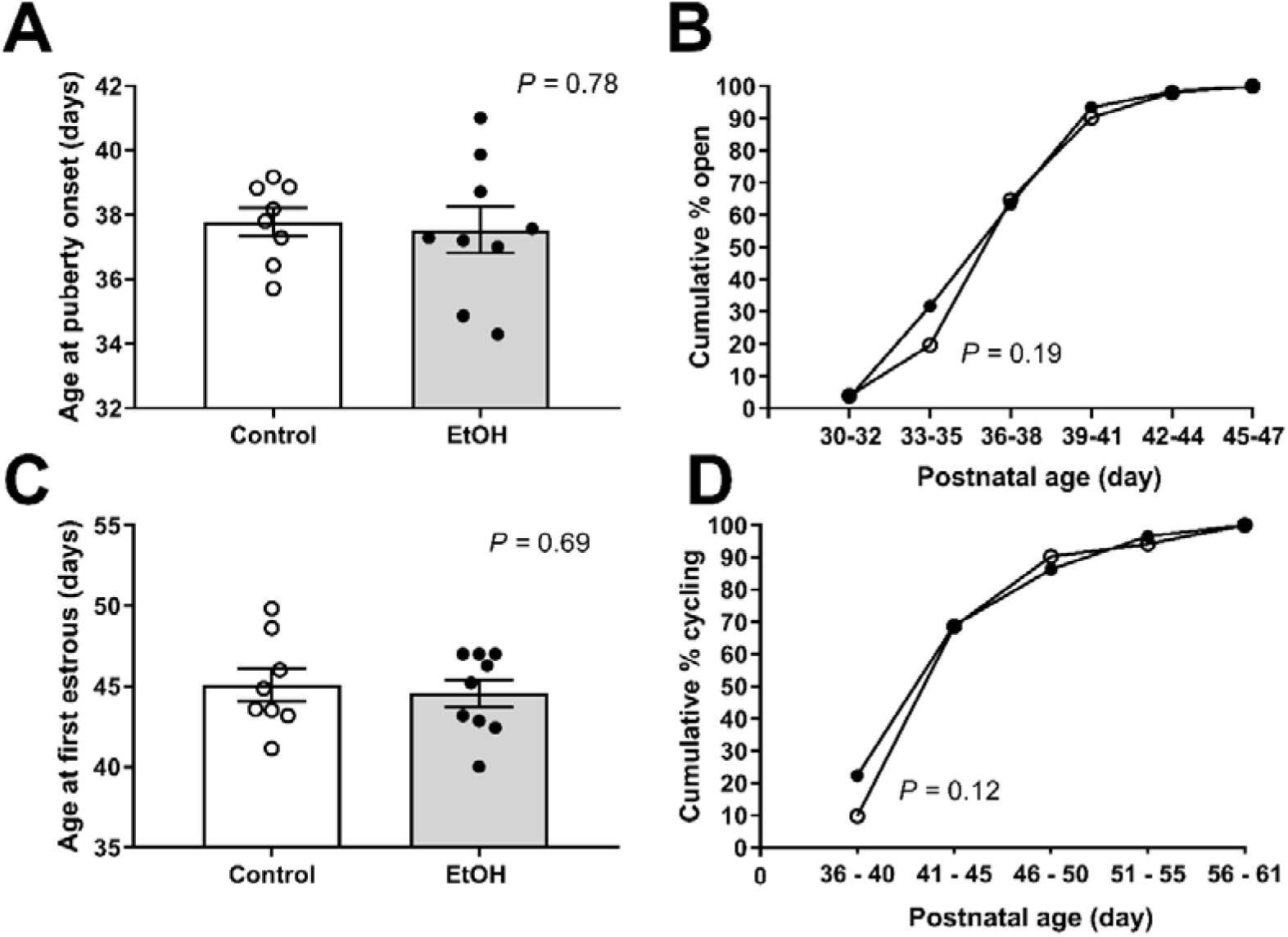
Moderate, acute prenatal alcohol exposure did not alter puberty onset in adolescent female rat offspring. Pregnant dams were treated with ethanol (EtOH, 1 g/kg body weight) or saline (control) at embryonic day (E) 13.5 and 14.5. (A) Age at puberty onset (as defined by vaginal opening). (B) Cumulative percentage of female offspring exhibiting puberty onset with increasing age. (C) Age at first estrous, as determined by vaginal electrical impedance >4kΩ. (D) Cumulative percentage of female offspring with estrous cycles with increasing age. For A) and C), open bars/circles indicate control offspring (litter averages from *n* = 8 litters; 5-7 females per litter); grey bars/solid circles indicate EtOH offspring (litter averages from n = *9* litters); all data are presented as mean ± SEM. Data analysed using unpaired t-tests. For B) and D), open circles indicate control offspring; solid circles indicate EtOH offspring; all data shown as % of all offspring across litters (number of litters as for A) and B)). Data compared at postnatal (PN) 33-35 in B) and at PN36-40 in D) using Fisher’s Exact test. Significance for all tests was set at *P*<0.05.

### Fertility parameters in adult offspring

PAE did not impact estrous cycle regularity in adult offspring. A typical 4-day cycle, as shown by the peaks in vaginal electrical impedance, is shown in Figure 4A. The number of estrous cycles across the 21-day monitoring period (Figure 4B) and the estrous cycle length (Figure 4C) did not differ between the Control and EtOH-exposed offspring. Plasma estradiol levels in pro-estrous animals also did not differ (Figure 4D).

**Figure 4.**
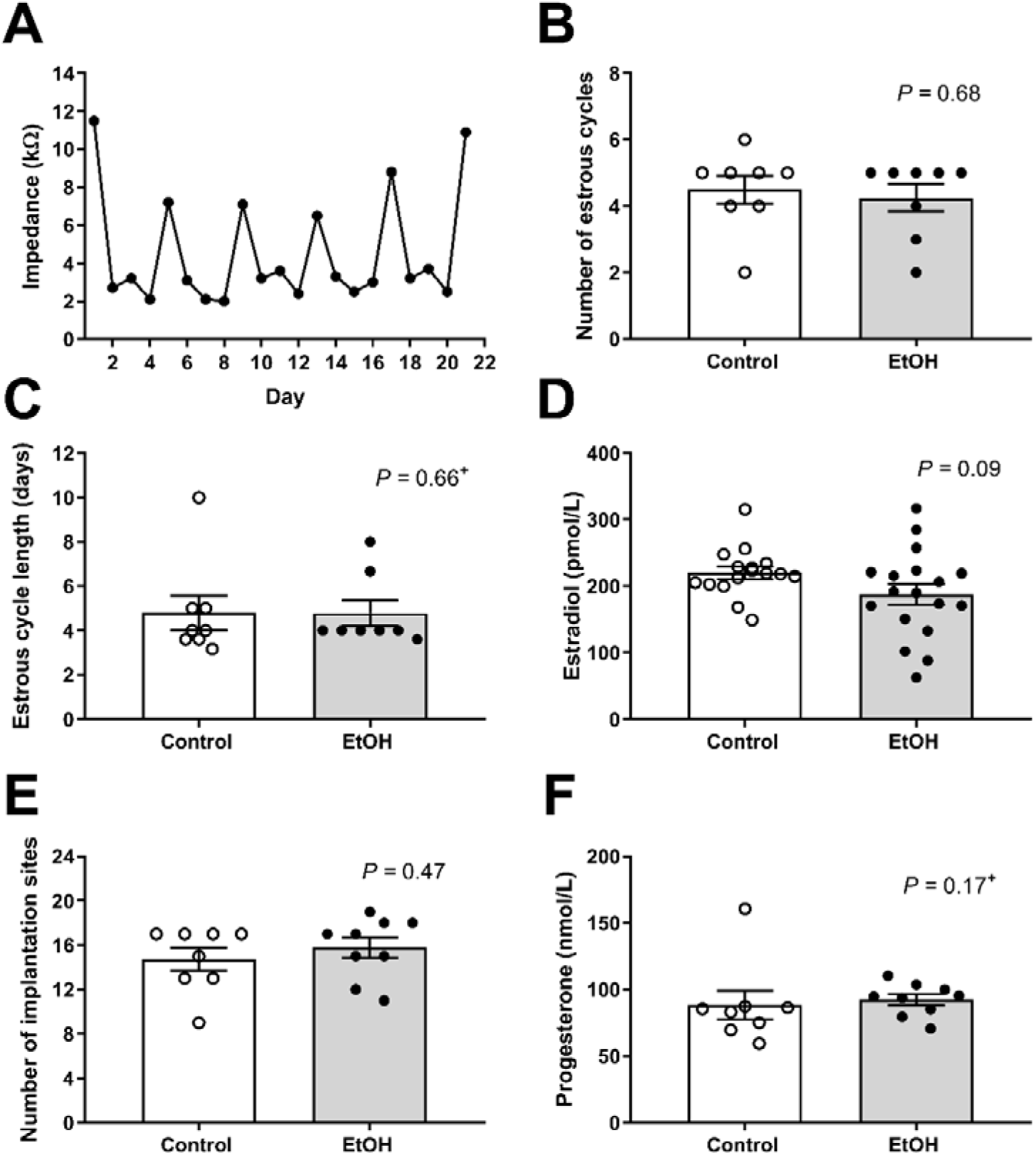
Moderate, acute prenatal alcohol exposure did not alter reproductive outcomes in adult female rat offspring. Pregnant dams were treated with ethanol (EtOH, 1 g/kg body weight) or saline (control) at embryonic day (E) 13.5 and 14.5. (A) An example of vaginal impedance readings over a 21-day monitoring period, indicating regular estrous cycles. Peaks >4kΩ indicate estrous. (B) Number of estrous cycles measured over the 21-day monitoring period. (C) Average length of estrous cycles (per female) measured over the 21-day monitoring period. For B) and C), *n* = 8 per group (1 female from each litter). (D) Plasma estradiol in 6-month old female offspring during estrous (vaginal impedance >4kΩ); *n* = 16 (control) or 18 (EtOH) per group (2 females from each litter). E) Number of implantation sites per mated female offspring at embryonic day (E) 9-11. (F) Plasma progesterone levels of mated female offspring at E9-11. For E) to F), *n* = 8 (control) or 9 (EtOH) per group (1 female from each litter). For B) - F), open bars/circles indicate control offspring; grey bars/solid circles indicate EtOH offspring; all data are presented as mean ± SEM. Data were analysed using an unpaired t-test (parametric data) or non-parametric Mann-Whitney rank sum test (indicated by ^+^) to determine significant differences between groups, with significance determined at *P*<0.05.

Following mating with a proven stud male at pro-estrous, all animals became pregnant within 5h post-mating, as shown by the presence of a seminal plug. Implantation rates at E9-11were similar between the two groups, with dissected uteri containing an average of 15 implantation sites (Figure 4E). PAE also did not impact the fetal resorption rate, with uteri of both Control and EtOH offspring containing an average of 2 resorption sites (data not shown). Plasma progesterone levels at this stage of pregnancy were also not different between groups (Figure 4F).

## Discussion

While consumption of alcohol in women has been linked to reduced ovarian reserve and impaired reproductive success, the effect of PAE on these reproductive outcomes in female offspring is unknown. To our knowledge, this is the first study to conduct a comprehensive investigation of reproductive parameters across the life course in female offspring following a relatively modest, acute dose of alcohol during pregnancy. Our results suggest that this model is not detrimental to the follicle pool established immediately after birth, does not change the age of puberty onset and the initiation of estrous, and does not affect estrous cycles and fertility in adult female offspring. These results provide some reassurance for women who may have consumed a low level of alcohol during pregnancy.

Previous preclinical studies investigating potential effects of PAE on female reproductive outcomes in offspring have typically used high doses of alcohol (35-36% EtOH-derived calories) in a Lieber-deCarli style liquid diet (see Akison, et al. (2019) for review), resulting in BACs of ∼100-150 mg/dl (0.10-0.15%) (Elton, et al. 2002). This is equivalent to 3-5 standard drinks consumed in two hours by an average weight woman (Leeman, et al. 2010). We report here that the average BAC across all EtOH-treated dams was ∼42 mg/dl (0.04%), slightly less than the 0.05% reported in Nguyen, et al. (2019) due to slightly lower BACs in the additional six treated dams used in this study for neonatal offspring studies. This was despite a consistent dosage, normalised to body weight, and highlights the individual variability in response to EtOH treatment and the importance of monitoring and reporting this response via BAC levels. Thus, the much lower BAC level compared to previous studies may explain the lack of a significant reproductive phenotype in PAE offspring. Given the paucity of studies examining the effect of PAE on ovarian reserve and later fertility, it remains to be seen if higher doses of alcohol can impact these reproductive parameters. Of note, examples of previous preclinical studies showing significant impacts on ovarian reserve associated with maternal smoke exposure (Camlin, et al. 2016) and undernutrition (Bernal, et al. 2010, Winship, et al. 2018) used much more severe exposures, with the equivalent of 12 cigarettes/75 min exposure twice daily in the former study and a 50% or greater reduction in calories or protein compared to the control intake in the latter studies.

The timing of exposure corresponded with an important stage in prenatal ovarian development in the rat, when germ cells cysts or ‘nests’ form, and the first wave of apoptosis occurs (Sarraj and Drummond 2012). We used an acute exposure of only two doses on two consecutive days of pregnancy, designed to mimic a low level of exposure over a weekend or on a special occasion. This is the first time that reproductive outcomes following such a short exposure have been examined, with previous preclinical studies typically treating throughout pregnancy or from mid-gestation to birth (see Akison, et al. (2019) for review). Again, this may explain the lack of a significant reproductive phenotype, and further studies are needed to determine if establishment of ovarian reserve is altered if exposure to alcohol continues through late gestation until birth, when primordial follicles are forming. Indeed, previous examples showing significant effects of maternal insults on offspring ovarian reserve were conducted throughout pregnancy and/or lactation (Bernal, et al. 2010, Camlin, et al. 2016), with one recent study finding effects of a maternal low-protein diet also including the preconception period (Winship, et al. 2018).

Previous preclinical studies have reported that more extensive, higher dose PAE is associated with delays in puberty onset in six out of eight studies (see Akison, et al. (2019) for review). Most studies reported this as the cumulative percentage of females displaying vaginal opening over time. Two studies in different rat strains showed that 100% of controls had achieved puberty by PN40-42, while only 73% of EtOH-exposed offspring achieved this developmental milestone by this age, with some animals delayed to as late as PN46 (Esquifino, et al. 1986, McGivern and Yellon 1992). Aside from puberty onset, three preclinical studies, all in rats, have previously examined estrous cyclicity using vaginal cytology, with only one study reporting an increased incidence of acyclic females which emerged with increasing age (i.e. at 6 and 12 months but not at 2 months; McGivern, et al. (1995)). The other two studies examined estrous cycles at 3.5-4 months of age and found no differences (Hard, et al. 1984, Lan, et al. 2009). This suggests that PAE may shorten the reproductive lifespan, which is only apparent when examining older females. In our study, we examined estrous cycles in female offspring at 6 months of age, an age of peak reproductive capacity, and found no differences. Rats typically undergo reproductive senescence (equivalent to menopause in women) at around 12 months of age, with subfertility at around 8-10 months of age (see Cruz, et al. (2017) for review). Thus, future studies examining PAE in aged animals may be of benefit. This subfertile period typically occurs in humans at 37.5-51 years of age in humans, culminating in menopause at ∼51 years of age (te Velde 1998).

Few studies (*n* = 4) have previously examined hormone levels in female offspring following PAE (see Akison, et al. (2019) for review), with only one study in 10 week-old rats reporting increased circulating estradiol levels at pro-estrous compared to controls (Polanco, et al. 2010). We found no effect of PAE on plasma estradiol at the same stage of the estrous cycle at 6 months of age, nor on plasma progesterone levels during early pregnancy. There were also no direct effects of ethanol on progesterone levels in late pregnancy in treated dams, which is contrary to reports of reduced progesterone concentrations in response to alcohol exposure, albeit in pre-menopausal, non-pregnant women (Gill 2000, Rachdaoui and Sarkar 2017).

We measured expression of a subset of potential factors involved in regulating primordial follicle numbers to determine if these were affected by PAE. Numbers of these non-growing follicles are controlled by a balance of factors that regulate activation and recruitment to the growth phase and maintain quiescence via extrinsic (e.g. *Amh, Inha*) (Durlinger, et al. 2002, Myers, et al. 2009, Pangas 2012) and intrinsic (e.g. *Pten, Cxcr4, Cxcl12, Stk11/Lkb1*) (Holt, et al. 2006, Jiang, et al. 2016, Reddy, et al. 2008, Reddy, et al. 2010, Wear, et al. 2016) factors; and regulate depletion via pro- and anti-apoptotic factors (e.g. *Bax, Bak1, Bcl2, Bcl2l1, Bcl2l11, Puma/Bbc3*) (Flaws, et al. 2001, Hutt 2015, Liew, et al. 2014, Liew, et al. 2016, Myers, et al. 2014). While many other factors are emerging as being important, we chose this subset to encompass these three major fates of primordial follicles – activation, quiescence and apoptosis. Consistent with our follicle count data, we found no significant difference in any of these factors, at least at the mRNA level, in neonatal ovaries exposed to PAE compared to controls. We chose the early (PN3) and late (PN10) neonatal periods, as these correspond with peak numbers of primordial follicles and a period of active recruitment to the growing follicle pool in the rat (Picut, et al. 2015). We did not look for any differences in ovarian reserve in young or aged adult ovaries, given no evidence for differences in the starting population of primordial follicles in neonates, no differences in molecular factors regulating follicle numbers and no differences in the number of implantations in mated adults, which is reflective of the number of ovulated follicles in that cycle.

Although we found no effects of PAE in this study, we emphasise that this same model did produce sex-specific metabolic effects in offspring (Nguyen, et al. 2019), with males at 6 months of age, litter-mates of the adult females reported here, showing evidence of insulin resistance. Female offspring were also examined but showed no evidence of metabolic dysfunction. This suggests that this model of PAE can result in sex-specific programming of long-term health. Therefore, it would be interesting for future studies to examine the impact of this low-level PAE model on male reproductive parameters, especially as there is some evidence that higher doses of alcohol during pregnancy can affect testis development and long-term reproductive performance (Lan, et al. 2013, Udani, et al. 1985). Also, our previous work in a model of PAE administered at a similar stage of gestation, but resulting in a higher BAC, impaired renal development and resulted in a low nephron endowment with offspring developing hypertension and renal dysfunction (Gray, et al. 2010). This suggests alcohol consumption at this stage of gestation can cause significant long-term health problems but the amount consumed and sex of the fetus plays an important role in determining outcomes.

### Conclusions

Given that women are increasingly delaying child-bearing, it is imperative that nutritional or environmental insults during pregnancy, that could impact on the establishment of the primordial follicle pool and hence the reproductive lifespan, are identified. Results from this preclinical study suggest that a moderate, episodic exposure to alcohol during pregnancy does not impact on ovarian reserve and subsequent fertility in adulthood. However, the previously reported metabolic deficits in male offspring from the same litters indicates that abstaining from alcohol consumption during pregnancy is the safest option for long-term offspring health.

## Author contribution statement

LKA conceived the study, analysed and finalised data, wrote the first draft of the manuscript and finalised the final version for submission. EKM and SES performed experiments, tissue preparation, assisted with data analysis and contributed to a first draft of the manuscript. SHL and KJH provided training and intellectual input for stereology. KMM provided intellectual input on study design and analysis and contributed to early drafts of the manuscript. All authors contributed to editing early drafts of the manuscript.

## Acknowledgements

The authors would like to acknowledge Natasha Steiger (Animal Endocrinology Lab, School of Biomedical Sciences, University of Queensland) for analysis of plasma hormones; Dave Sterne and Barb Arnts (University of Queensland Biological Resources) for animal treatments and husbandry; Erica Mu (School of Biomedical Sciences Histology Facility, University of Queensland) for serial sectioning of neonatal ovaries; Shaun Walters (School of Biomedical Sciences Imaging Facility, University of Queensland) for technical assistance with stereology; Tam Nguyen (School of Biomedical Sciences, University of Queensland) for assistance with animal work; and Jacobus Ungerer (Pathology Queensland, Queensland Health) for analysis of BAC.

